# Neuropathological Correlates of Apathy Progression in Alzheimer’s Disease and Related Dementias: A Longitudinal NACC Cohort Study

**DOI:** 10.1101/2025.11.06.686966

**Authors:** Yumeng Qi, Terry E. Goldberg, Hyun Kim, D. P. Devanand, Seonjoo Lee

## Abstract

**Objective:** Apathy is a prevalent and disabling symptom in Alzheimer’s disease and related dementias, yet its progression across neuropathological subtypes remains incompletely understood. This cohort study investigates longitudinal changes in apathy and their associations with major neuropathologies using data from the National Alzheimer’s Coordinating Center.

**Methods:** We analyzed 1,488 participants with autopsy-confirmed neuropathology and at least two caregiver-reported NPI-Q assessments. Generalized linear mixed models were used to assess associations between apathy and six neuropathologies—Alzheimer’s disease (AD), Lewy body disease (LBD), frontotemporal lobar degeneration (FTLD), hippocampal sclerosis (HS), cerebrovascular disease (CVD), and cerebral amyloid angiopathy (CAA). with time modeled as years to death and including interaction terms. Sex-stratified analyses were also conducted. All models were adjusted for age at death, sex, and NPI-Q total score excluding apathy.

**Results:** Apathy prevalence increased over time across all pathology groups. FTLD (OR = 2.11, 95% CI: 1.32–3.39) and HS (OR = 2.23, 95% CI: 1.38–3.60) were consistently associated with higher odds of apathy throughout the disease course. No significant interaction effect was observed in any of the neuropathologies. In sex-stratified analyses, FTLD (OR=2.58, 95% CI 1.40-4.77), HS (OR=2.48, 95% CI 1.29-4.77), and LBD (OR=1.64, 95% CI 1.03-2.61) were significantly associated with apathy in males, while only HS (OR=2.18, 95% CI 1.06-4.47) remained significant in females.

**Interpretation:** Apathy severity varied by neuropathologies but progressed similarly over time. The elevated burden in FTLD and HS, particularly among males, underscores the importance of stratified approaches to early detection and intervention targeting apathy in ADRDs.

## Introduction

Apathy is one of the most prevalent and persistent neuropsychiatric symptoms (NPS) in individuals with Alzheimer’s disease and related dementias (ADRD), including Alzheimer’s disease (AD), Lewy body disease (LBD), cerebral amyloid angiopathy (CAA), cerebrovascular disease (CVD), frontotemporal lobar degeneration (FTLD), and hippocampal sclerosis (HS).^1–4^ Defined as a marked reduction in goal-directed behavior not attributable to primary cognitive, emotional, or motor impairments, apathy encompasses diminished motivation, initiative, and emotional responsiveness.^5–7^ It has been linked with worse functional outcomes, increased caregiver burden, diminished quality of life, and accelerated cognitive decline. ^8–11^

While prior studies have demonstrated a longitudinal association between apathy and tau and/or amyloid burden among AD patients using CSF biomarkers and PET imaging,^12–14^ such approaches fall short of the diagnostic certainty afforded by postmortem neuropathological examinations. Existing postmortem neuropathological data have primarily relied on cross-sectional designs or aggregated measures, limiting the ability to capture the dynamic progression of apathy across the course of ADRD. ^15,16^

In this study, we analyzed the longitudinal clinical and neuropathological data from the National Alzheimer’s Coordinating Center (NACC) to examine the temporal progression of apathy and its associations with distinct ADRD neuropathologies.^17^ Using years to death as a proxy for disease stage, we evaluated pathology-specific contributions to apathy over time. We further explored sex differences in these associations. Our findings aim to elucidate the neuropathological mechanisms underlying apathy in ADRD and inform more tailored approaches to early detection and intervention.

## Methods

### Standard Protocol Approvals, Registrations, and Patient Consents

Neuropsychiatric and neuropathologic data were obtained from the NACC, which is the data repository for past and present Alzheimer’s Disease Research Centers (ADRCs) funded by the National Institute on Aging.^17^ ADRCs obtained written informed consent from each included participant (or guardians of participants) in the study, and each institution maintained its own separate Institutional Review Board review process.

### Study Sample

At 39 US Alzheimer Disease Center sites, autopsies were conducted locally using the NACC Coding Guidebook protocol (January 2014) for uniform collection and ratings of neuropathological data. Institutional review board approval for clinical and autopsy procedures were obtained locally.

Neuropathological data collected between January 2012 and January 2018 were compiled into the publicly available NACC v.10 database.^17^ Cases with neuropathological diagnoses of brain injury, central nervous system neoplasm, Down syndrome, Huntington’s disease, or prion disease were excluded. To ensure stable estimation in longitudinal modeling, we included only individuals with at least two the caregiver-reported Neuropsychiatric Inventory Questionnaire (NPI-Q) assessments.

According to the NACC protocol, to make a neuropathological diagnosis, semiquantitative neuropathological assessments were performed using immunohistochemistry, histochemistry, microscopic visualization, or visual inspection, with region-specific categorization. AD neuropathologic change was evaluated with an ‘ABC’ score: Aβ/amyloid plaques(A), NFT stage (B), and neuritic plaque score (C). The combination of A, B, and C scores is designated as “Not,” “Low,” “Intermediate,” or “High” AD neuropathologic change. Intermediate or High AD neuropathologic change is labeled as presence^18^ Lewy body pathology was determined by alpha-synuclein immunohistochemistry (IHC).^18,19^ FTLD was defined as present or absent based on any of the following diagnoses: FTLD with tau pathology or other tauopathy, FTLD with TDP-43 pathology,^20^ Other FTLD subtype.^18,21,22^ CAA was defined as mild to severe global burden, corresponding to scattered to widespread vascular amyloid deposition.^23,24^ CVD was defined as multiple microinfarcts or infarcts or lacunes.Cases with severe neuronal loss and gliosis in CA1 and/or subiculum were included as hippocampal sclerosis.^18^

### Apathy

Apathy was assessed using the NPI-Q, a widely used informant-based took that evaluates 12 neuropsychiatric symptoms, including apathy.^25^ Informants were asked to indicate whether each symptom had been present during the month prior to the assessment. If present, symptom severity was further rated on a 3-point scale, 1 (Mild), 2 (Moderate), and 3 (Severe). In this study, apathy was primarily analyzed as a binary outcome—present versus absent—while severity ratings were examined in supplementary analyses.

### Statistical Analyses

All analyses were conducted using R (version 4.4.1). Means and standard deviations (SDs) were reported for continuous variables, and frequencies with percentages were reported for categorical variables. Group comparisons were performed using two-sample *t*-tests for continuous variables and chi-squared tests for categorial variables.

To examine the longitudinal association between neuropathological subtype and apathy progression, we used GLMM with a logit link function and binomial distribution. Time was modeled as a continuous variable, defined by years to death (YTD), and interaction terms between each pathology and YTD were included to assess differential trajectories. In addition to the primary binary outcome models, we conducted secondary analyses treating apathy severity as an ordinal outcome using continuation-ratio models. These models accounted for the ordered nature of severity ratings (absent, mild, moderate, severe) and allowed us to examine whether specific neuropathological subtypes were associated with greater severity of apathy across time.

A series of models were estimated to examine the association between neuropathology and apathy. Models 1 and 2 evaluated each neuropathological subtype independently, without adjusting for other pathologies. In Model 1, the reference group comprised individuals without the specific target pathology. As such they were a mixture of all other neuropathology. In Model 2, the reference group was the individuals with no known neuropathology (NoNP). Models 3 and 4 further assessed the independent effects of all neuropathologies simultaneously on apathy progression, with Model 4 additionally adjusted for no-known pathology. In addition, we examined the most frequent comorbidity combinations. For these analyses, Models 5 and 6 were specified in parallel with Models 1 and 2: Model 5 used all other groups as the reference, and Model 6 used the NoNP group as the reference. This allowed us to assess whether common overlapping pathologies were associated with distinct trajectories of apathy.

Sex-stratified analyses were conducted to explore sex differences in neuropathology. All models were adjusted for sex, age at death, and NPI-Q total score excluding the apathy subscale. To account for the clinical overlap and potential confounding effects of depression, a sensitivity analysis was conducted with additional adjustment for depression status. Odds ratios (ORs) and 95% confidence intervals (CIs) were reported. Statistical significance was determined at thresholds of *p* ≤ 0.05. We performed multiple comparisons correction controlling for false discovery rate.

## Data Availability

Anonymized data not published within this article will be made available by request from any qualified investigator.

## Results

### Participants

A total of 1488 participants with available neuropsychiatric and neuropathological data were included in the analysis (See Table 1). The mean age at death was 80.91 (SD 10.67) years. Of the included individuals, 817 (54.9%) were male, and 1409 (94.7%) identified as Caucasian/White. The mean years of education across all participants was 15.51 (SD 3.10) and the average number of annual clinical visits was 5.28 (SD 2.37).

**Table 1.** Demographic and Clinical Characteristics by Neuropathological Diagnosis.

| Characteristic | Total<br>(N=1488) | AD<br>(N=1049) | LBD<br>(N=556) | CAA<br>(N=934) | CVD<br>(N=475) | FTLD<br>(N=296) | HS<br>(N=223) | NoNP<br>(N=93) |
| --- | --- | --- | --- | --- | --- | --- | --- | --- |
| Age at initial visit, mean (SD), y | 74.34<br>(10.09) | 74.65<br>(9.77) | 73.53<br>(9.41) | 74.59<br>(9.98) | 78.17<br>(8.78) | 71.54<br>(10.82) | 75.96<br>(9.39) | 72.28<br>(11.13) |
| Age at last visit, mean (SD), y | 79.30<br>(10.85) | 79.69<br>(10.38) | 78.28<br>(10.01) | 79.65<br>(10.59) | 83.21<br>(9.36) | 75.79<br>(11.81) | 81.09<br>(10.01) | 77.12<br>(12.44) |
| Age at death, mean (SD), y | 80.91<br>(10.67) | 81.36<br>(10.20) | 80.12<br>(9.78) | 81.33<br>(10.38) | 84.98<br>(9.14) | 77.36<br>(11.71) | 82.96<br>(9.74) | 78.35<br>(12.49) |
| Sex (Male), n (%) | 817<br>(54.9%) | 551<br>(52.5%) | 330<br>(59.4%) | 509<br>(54.5%) | 239<br>(50.3%) | 171<br>(57.8%) | 112<br>(50.2%) | 67<br>(72.0%) |
| Education, n (%) | 15.51<br>(3.10) | 15.44<br>(3.15) | 15.73<br>(3.07) | 15.43<br>(3.16) | 15.32<br>(3.16) | 15.78<br>(2.83) | 15.51<br>(3.18) | 15.90<br>(3.00) |
| Race (Caucasian /White), n (%) | 1409<br>(94.7%) | 994<br>(94.8%) | 531<br>(95.5%) | 883<br>(94.5%) | 451<br>(94.9%) | 286<br>(96.6%) | 209<br>(93.7%) | 88<br>(94.6%) |
| Number of visits, mean (SD) | 5.28<br>(2.37) | 5.31<br>(2.37) | 5.10<br>(2.34) | 5.31<br>(2.36) | 5.32<br>(2.37) | 4.64<br>(2.24) | 5.42<br>(2.34) | 5.30<br>(2.51) |
| Apathy at last visit, n (%) | 711<br>(47.8%) | 515<br>(49.1%) | 286<br>(51.4%) | 446<br>(47.8%) | 203<br>(42.7%) | 170<br>(57.4%) | 121<br>(54.3%) | 34<br>(36.6%) |
| Total NPI at last visit, mean (SD) | 5.81<br>(5.59) | 6.08<br>(5.69) | 6.19<br>(5.66) | 5.96<br>(5.64) | 5.33<br>(5.27) | 6.31<br>(5.39) | 5.92<br>(4.83) | 5.05<br>(5.99) |

**Notes:** Means and SDs were reported for continuous variables, while frequencies and percentages were used for categorical variables. Group comparisons were performed using t-tests and chi-squared test as appropriate. Abbreviations: AD, Alzheimer disease (defined as intermediate or high ABC criteria); CAA, cerebral amyloid angiopathy; CVD, cerebrovascular disease; FTLN, frontotemporal lobar degeneration; HS, hippocampal sclerosis; LBD, Lewy body disease; NoNP, no known pathology.

Figure 1A shows the distribution of participants by neuropathologies, with AD and CAA dominating the cohort (70.5% and 62.8%, respectively). A small proportion of the sample (6.2%) had no known pathology. In the study cohort, many participants presented with co-occurring neuropathologies, the largest comorbidity group was AD with CAA (n = 238). This was followed by AD with LBD and CAA (n = 201) and AD with CAA and CVD (n = 120, see eTable 1). Among the six neuropathologies, FTLD had the highest rate of missingness (334, 22.4%, see eFigure 1).

**Figure 1.**
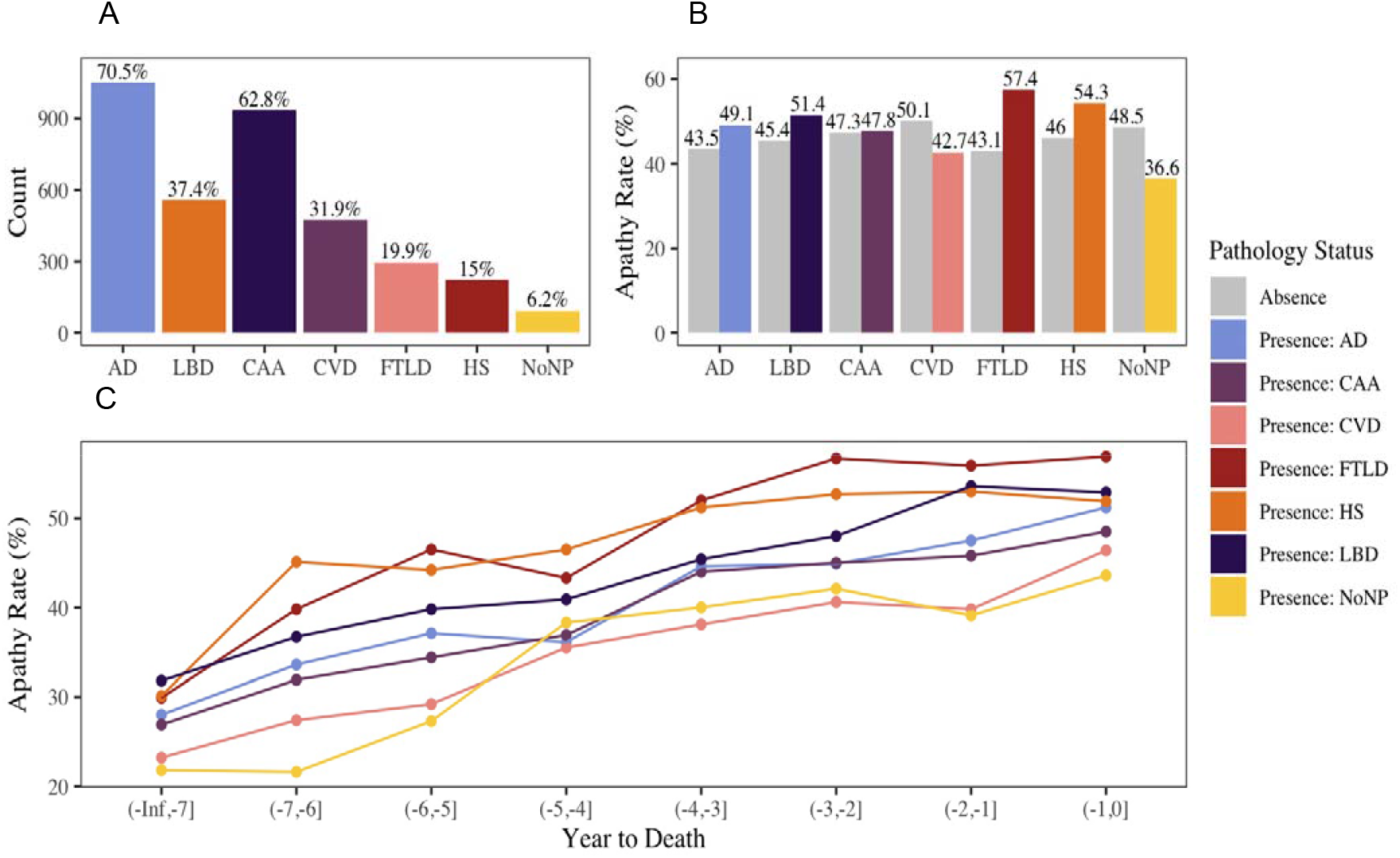
Prevalence of neuropathology and apathy across neuropathology. (A) Distribution of neuropathology in the sample (presence percentage annotated above each bar). (B) Apathy prevalence (%) stratified by the presence or absence of each neuropathology. (C) Longitudinal apathy trajectories over time (years to death) across neuropathology groups, showing a general increase in apathy prevalence approaching death. Bar and line colors indicate the presence of each neuropathology (legend, right), with gray bars representing pathology absence. Abbreviations: AD, Alzheimer disease (defined as intermediate or high ABC criteria); CAA, cerebral amyloid angiopathy; CVD, cerebrovascular disease; FTLD, frontotemporal lobar degeneration; HS, hippocampal sclerosis; LBD, Lewy body disease; NoNP, no known pathology.

At the last visit, the overall mean NPI-Q total score was 5.81 (SD 5.59). Apathy as assessed by the NPI-Q was observed in 711 participants (47.8%) overall, with the highest prevalence seen in individuals with FTLD (57.4%). Figure 1B showed apathy prevalence at last visit stratified by the presence or absence of each pathology. In general, individuals with a given pathology showed higher apathy prevalence than those without, particularly for FTLD (57.4% vs 43.1%), HS (54.3% vs 46.0%), and LBD (51.4% vs 45.4%). However, this pattern did not hold for CVD (presence vs absence, 42.7% vs 50.1%). Figure 1C showed that apathy prevalences increased over time across all neuropathological groups, with FTLD and HS consistently showing the highest rates across time bins. While baseline prevalence differed slightly across the groups, most groups demonstrate an overall increase of about 20 to 25 percentage points from early to late time points.

### Longitudinal Analysis

Table 2 summarizes the longitudinal associations between apathy and each neuropathology of interest, without adjusting for the presence of other neuropathologies. In Model 1, compared to the absence of the target pathology, FTLD (OR = 2.11, 95% CI: 1.32–3.39, *p* < 0.01) and HS (OR = 2.23, 95% CI: 1.38–3.60, *p* < 0.001) were associated with greater odds of apathy. Years to death was also associated with apathy across all pathology groups (ORs ranging from 1.12 to 1.18, *p* < 0.001), indicating that apathy became more prevalent over time. No significant interactions between neuropathology and time were observed. In Model 2, compared to the NoNP group, FTLD (OR = 2.91, 95% CI: 1.27–6.63, *p* < 0.05), HS (OR = 3.87, 95% CI: 1.69–8.86, *p* < 0.001), and LBD (OR = 2.45, 95% CI: 1.15–5.21, *p* < 0.05) were associated with greater odds of apathy. YTD was not significantly associated with apathy in this model, and no significant neuropathology-by-time interaction effects were detected.

**Table 2.**
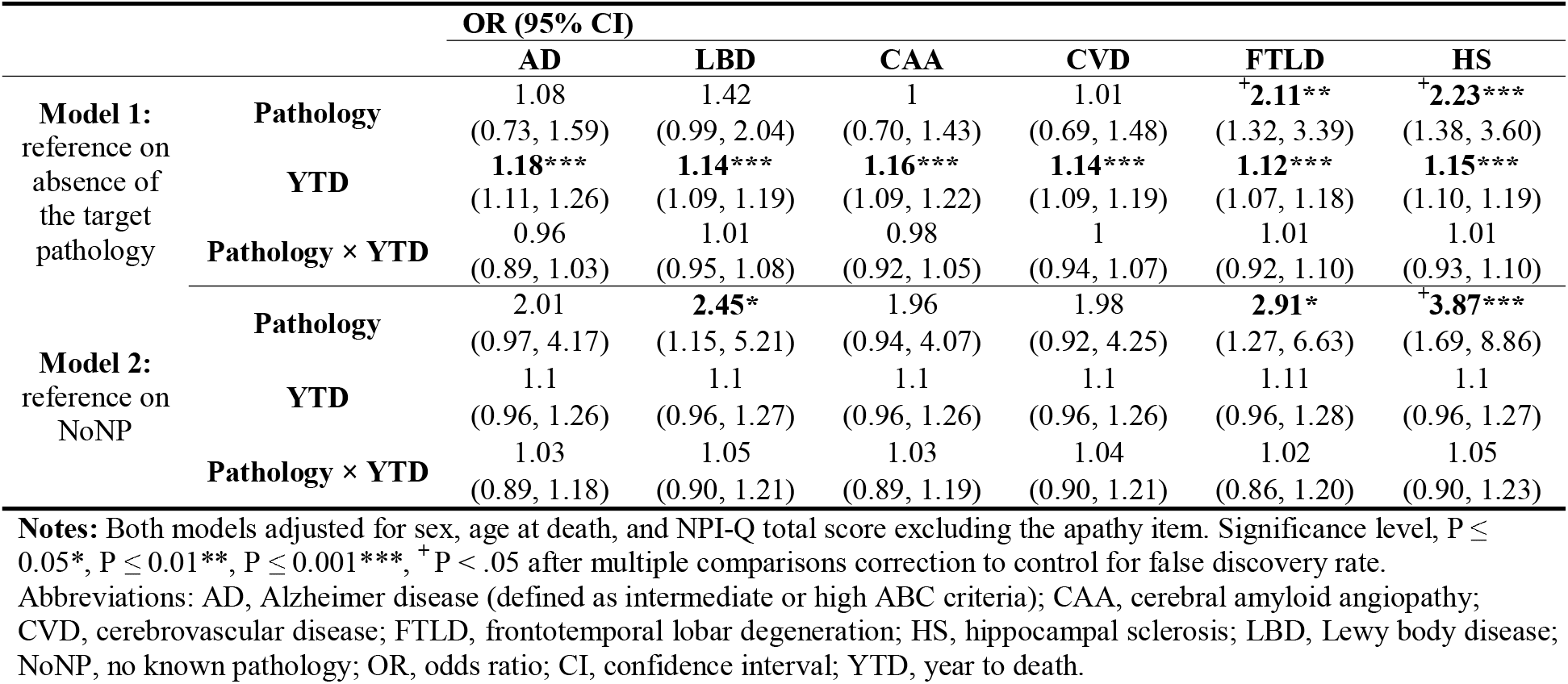
Association Between Apathy and Each Neuropathology Without Controlling for Other Pathologies.

Sex-stratified analysis were largely consistent among males: in Model 1, FTLD (OR = 2.58, 95% CI: 1.40–4.77, *p* < 0.01), HS (OR = 2.48, 95% CI: 1.29–4.77, *p* < 0.01), and LBD (OR = 1.64, 95% CI: 1.03–2.61, *p* < 0.05) were associated with greater odds of apathy when each were referenced on the absence of target neuropathology, and YTD remained a strong predictor across all neuropathologies. In Model 2, FTLD (OR = 3.63, 95% CI: 1.36–9.72, *p* < 0.01), HS (OR = 4.31, 95% CI: 1.56–11.94, *p* < 0.01), and LBD (OR = 2.67, 95% CI: 1.09–6.52, *p* < 0.05) were associated with apathy compared to the NoNP group. No significant neuropathology-by-time interactions were observed. Among female participants, in Model 1, HS was associated with apathy (OR = 2.18, 95% CI: 1.06–4.47, *p* < 0.05), and YTD remained significant across all pathologies. No interaction effects were found. In Model 2, none of the neuropathology showed a significant association with apathy (see eTable 2).

Figure 2 summarizes the odds ratios and 95% CI from both models across all participants, males, and females. Not unexpectedly, across most neuropathologies, Model 2 (comparisons across NoNP and neuropathology groups) yielded higher odds ratios than Model 1 (comparisons within neuropathologies, none vs. presence of neuropathology of interest). Within each model, male participants generally showed higher odds ratios compared to all participants and females. When treating apathy severity as an ordinal outcome using an ordinal GLMM, apathy severity increases in FTLD and HS, but no interaction effects were found (see eTable 3 and eTable 4).

**Figure 2.**
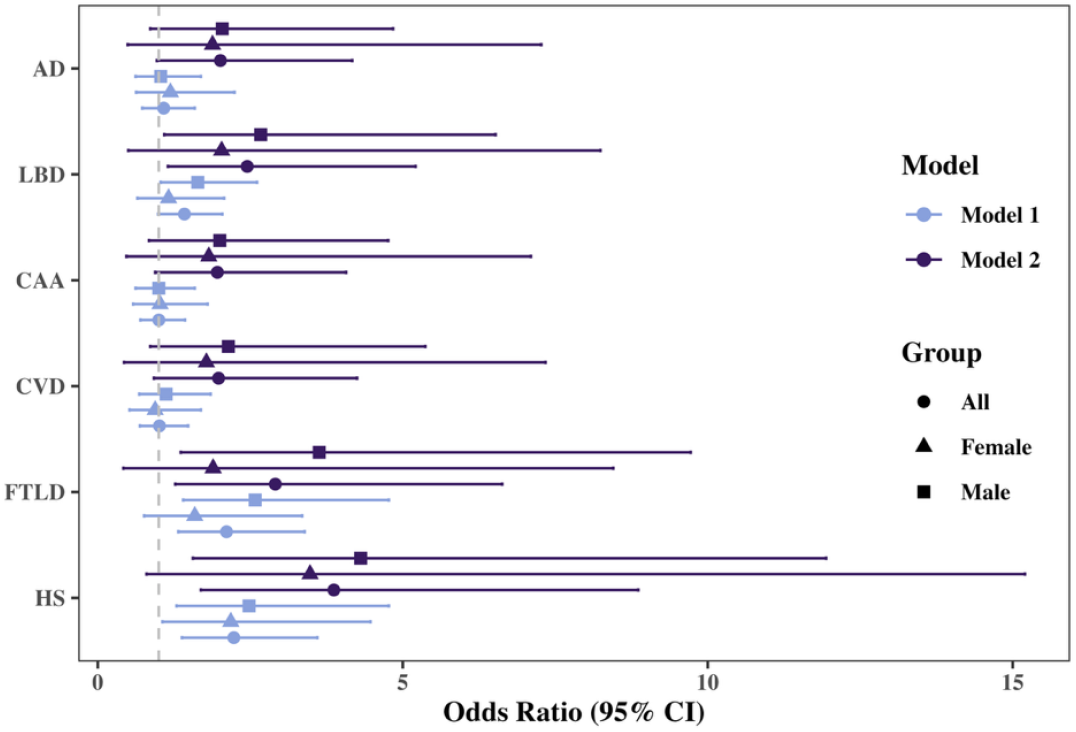
Forest Plot of OR and 95% CI. Points mark the OR estimates, and horizontal lines represent the corresponding 95% CIs. Color differentiates the model, model 1 referenced on absence of the target pathology, while model 2 referenced on NoNP. Shapes indicate participant groups, circles = all participants, triangles = females, squares = males. Abbreviations: AD, Alzheimer disease (defined as intermediate or high ABC criteria); CAA, cerebral amyloid angiopathy; CVD, cerebrovascular disease; FTLD, frontotemporal lobar degeneration; HS, hippocampal sclerosis; LBD, Lewy body disease; NoNP, no known pathology; OR, odds ratio; CI, confidence interval.

Table 3 presents the independent effects of neuropathologies on apathy progression, controlling for all included pathologies. In both Model 3 and Model 4, FTLD (Model 3 OR = 2.24, 95% CI: 1.30–3.85; Model 4 OR = 2.18, 95% CI: 1.27–3.76) and HS (Model 3 OR = 2.22, 95% CI: 1.26–3.92; Model 4 OR = 2.18, 95% CI: 1.23–3.85) were associated with greater odds of apathy. YTD remained a strong predictor of apathy in both models (p < 0.001), while no pathology-by-time interaction reached significance. (See sex-stratified analyses at eTable 5)

For comorbidity analysis, we focus on the three most frequent combinations—AD+CAA, AD+CAA+LBD, and AD+CAA+CVD. No significant associations were found between comorbidity and apathy (See eTable 6).

**Table 3.**
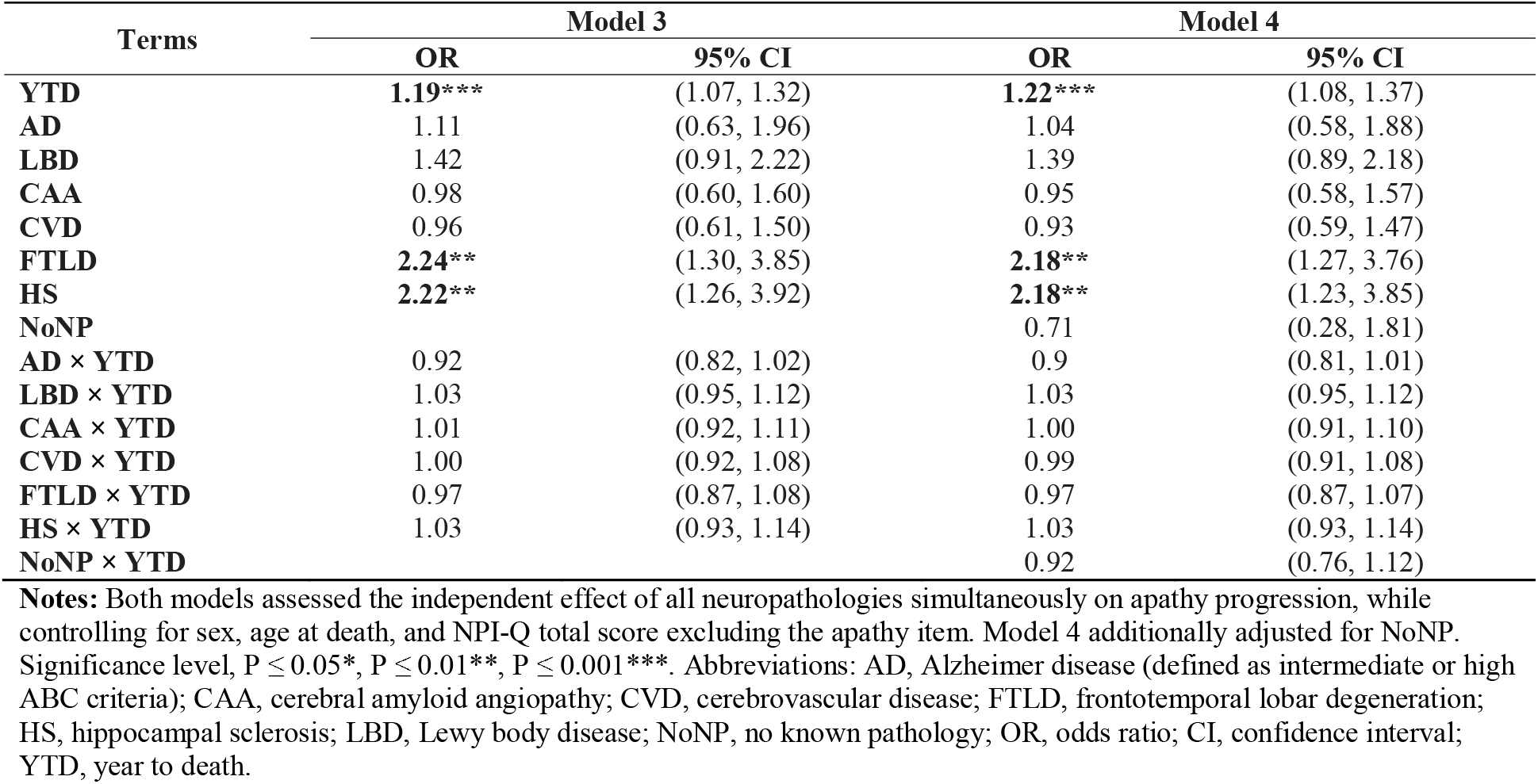
Independent Effect of Neuropathologies on Apathy Progression.

| Terms | Model 3 |  | Model 4 |  |
| --- | --- | --- | --- | --- |
|  | OR | 95% CI | OR | 95% CI |
| **YTD** | **1.19\*\*\*** | (1.07, 1.32) | **1.22\*\*\*** | (1.08, 1.37) |
| **AD** | 1.11 | (0.63, 1.96) | 1.04 | (0.58, 1.88) |
| **LBD** | 1.42 | (0.91, 2.22) | 1.39 | (0.89, 2.18) |
| **CAA** | 0.98 | (0.60, 1.60) | 0.95 | (0.58, 1.57) |
| **CVD** | 0.96 | (0.61, 1.50) | 0.93 | (0.59, 1.47) |
| **FTLD** | **2.24\*\*** | (1.30, 3.85) | **2.18\*\*** | (1.27, 3.76) |
| **HS** | **2.22\*\*** | (1.26, 3.92) | **2.18\*\*** | (1.23, 3.85) |
| **NoNP** |  |  | 0.71 | (0.28, 1.81) |
| **AD × YTD** | 0.92 | (0.82, 1.02) | 0.9 | (0.81, 1.01) |
| **LBD × YTD** | 1.03 | (0.95, 1.12) | 1.03 | (0.95, 1.12) |
| **CAA × YTD** | 1.01 | (0.92, 1.11) | 1.00 | (0.91, 1.10) |
| **CVD × YTD** | 1.00 | (0.92, 1.08) | 0.99 | (0.91, 1.08) |
| **FTLD × YTD** | 0.97 | (0.87, 1.08) | 0.97 | (0.87, 1.07) |
| **HS × YTD** | 1.03 | (0.93, 1.14) | 1.03 | (0.93, 1.14) |
| **NoNP × YTD** |  |  | 0.92 | (0.76, 1.12) |

**Notes:** Both models assessed the independent effect of all neuropathologies simultaneously on apathy progression, while controlling for sex, age at death, and NPI-Q total score excluding the apathy item. Model 4 additionally adjusted for NoNP. Significance level, $P \leq 0.05^*$ , $P \leq 0.01^{**}$ , $P \leq 0.001^{***}$ . Abbreviations: AD, Alzheimer disease (defined as intermediate or high ABC criteria); CAA, cerebral amyloid angiopathy; CVD, cerebrovascular disease; FTLD, frontotemporal lobar degeneration; HS, hippocampal sclerosis; LBD, Lewy body disease; NoNP, no known pathology; OR, odds ratio; CI, confidence interval; YTD, year to death.

### Sensitivity Analysis

Adjusting for depression did not alter our findings. The significant associations remained unchanged with slightly attenuated effect sizes. This consistency was observed across models with and without adjustment for other pathologies, as well as in sex-stratified analyses (See from eTable7 to eTable 10).

## Discussion

In this study, we examined longitudinal trajectories of apathy across six neuropathological groups of ADRD in a large, autopsy-confirmed cohort. Apathy rates increased over time in all neuropathology groups. Notably, FTLD and HS were consistently associated with higher odds of apathy throughout the disease course. However, we did not find significant interactions between time and neuropathology, suggesting that while certain pathologies increase the overall likelihood of apathy, they do not appear to influence the rate at which it progresses. Sex-stratified analyses revealed that FTLD, HS, and LBD were significantly associated with apathy in males, whereas in females, only HS remained significant. Notably, results remained nearly the same whether the reference group was “no known neuropathology” or “all neuropathologies” other than the target group.

Consistent with previous studies, our findings suggest that apathy symptoms persist throughout the course of AD.^26–28^ Moreover, we observed an overall increase in apathy prevalence over time across all neuropathologies, with no significant interaction between time and neuropathology status. This suggests that while the presence of certain neuropathology influences the baseline prevalence of apathy but does not significantly alter the rate of apathy progression as death approaches. As illustrated in Figure 1C, different neuropathologies appear to define distinct starting points for apathy prevalence, but the slopes of progression remain largely parallel across groups.

The lack of significant interactions between time and neuropathology suggests that apathy progression may be shaped by shared neurodegenerative mechanisms rather than pathology-specific effects. Once apathy emerges, its trajectory may reflect common disruptions to frontal-subcortical and limbic circuits, which are involved in motivation across multiple dementia subtypes.^29–31^ Additionally, though apathy has been regarded as an early indicator of AD,^32,33^ and some studies suggest distinct mechanisms may underlie apathy at early versus late disease stages^13,34^ our data primarily capture the last decade of life. Thus, earlier divergence in apathy trajectories may have gone undetected, highlighting the need for studies spanning earlier preclinical or clinical stages. Alternatively, interaction effects may be region-specific and thus not adequately captured by binary pathology classifications, underscoring the importance of considering disease subtypes and progression stages in future research.

The prevalence of apathy varied by neuropathology. AD and CAA were common in the cohort and frequently co-occurred, consistent with previous study.^35^ However, neither their presence alone nor in combination were associated with apathy. By contrast, FTLD and HS showed consistent strong associations. These findings align with prior research linking FTLD to prominent motivational and behavioral symptoms due to early involvement of medial frontal and cingulate regions—key nodes in apathy-related circuits.^36–38^ HS is associated with limbic system degeneration, affecting the hippocampus and surrounding medial temporal lobe structures, which may disrupt emotional regulation and motivational drive and result in observable apathy.^15,39^ The persistent and elevated odds of apathy in these groups across the disease course suggest that these neuropathologies may confer a higher and more stable vulnerability to apathy, independent of disease stage.

It is important to note that FTLD is a heterogeneous group of disorders with distinct underlying proteinopathies, primarily FTLD-TDP and FTLD-tau, each associated with different patterns of neurodegeneration.^21^ FTLD-TDP is often associated with anterior temporal and frontal lobe atrophy, while FTLD-tau subtypes (e.g., PSP, CBD, Pick’s disease) more commonly affect dorsolateral and medial prefrontal regions, basal ganglia, and brainstem structures.^40–42^ These distinct anatomical targets may disrupt different frontal-limbic and frontostriatal circuits implicated in goal-directed behavior and motivation, potentially leading to apathy via different neurobiological pathways. However, due to high missing rates in FTLD cases (22.4%) and limited subtype information in the current dataset, we were unable to disentangle the contributions of specific FTLD subtypes to apathy expression (See eTable 11 for FTLD subtype distribution).

Although HS and TDP-43 pathology are often reported to co-occur, particularly in older adults and individuals with AD,^43,44^ our dataset revealed surprisingly few overlapping cases. This limited overlap may reflect differences in sampling, cohort characteristics, or simply the missing rate in FTLD autopsy. Given the strong association observed between HS and apathy in our analyses, the limited co-occurrence with TDP-43 strengthens the interpretation that HS alone may be a key driver of motivational decline in late life.

Previous studies have reported sex differences in apathy prevalence among individuals with AD, with males often exhibiting higher rates of apathy than females.^45–47^ Consistent with this, our sex-stratified analyses further revealed that FTLD, HS, and LBD were significantly associated with greater odds of apathy in males, whereas in females, only HS remained significant, despite loss of power in this analysis. These findings suggest that the relationship between neuropathology and apathy may differ by sex, potentially due to differences in neurobiological vulnerability, hormonal influences, or sex-specific patterns of brain atrophy and functional decline.^48–50^ Understanding these distinctions could have important implications for developing personalized approaches to apathy screening and intervention in ADRD populations.

This study is subject to several limitations. First, the case series may be influenced by ascertainment bias, as a large proportion of participants met criteria for intermediate to high AD neuropathological change—a reflection of the NACC sites’ clinical focus on AD dementia. Second, a large majority of the sample was White, limiting generalizability to diverse populations. Third, the “no known neuropathology” group was cognitively heterogeneous, with over half meeting criteria for clinical dementia, highlighting the known discrepancies between clinical diagnoses and neuropathological confirmation. Fourth, data on FTLD were notably incomplete, with a missing rate of greater than 20%. Finally, apathy assessment via the NPI-Q relied on informant reports, which may be affected by recall bias or limited insight, potentially influencing symptom characterization. Future studies with enhanced apathy measures are warranted to clarify the relationship between ADRD neuropathology and the development of apathy.

## Conclusions

In this retrospective longitudinal study, apathy severity increased over time across six ADRD neuropathological groups, with FTLD and HS consistently associated with higher odds of apathy. These associations remained significant in males, with LBD additionally showing significant, whereas only HS remained significant in females. Notably, we did not observe significant interactions between time and apathy for any ADRD neuropathology, suggesting that while neuropathology influences overall apathy burden, it may not alter its progression rate. These findings highlight the role of shared neurodegenerative mechanisms in the development of neuropsychiatric symptoms and emphasize the clinical value of identifying apathy as a prognostic marker in dementia care.

## Acknowledgement statement

The NACC database is funded by NIA/NIH Grant U24 AG072122. NACC data are contributed by the NIA-funded ADRCs: P30 AG062429 (PI James Brewer, MD, PhD), P30 AG066468 (PI Oscar Lopez, MD), P30 AG062421 (PI Bradley Hyman, MD, PhD), P30 AG066509 (PI Thomas Grabowski, MD), P30 AG066514 (PI Mary Sano, PhD), P30 AG066530 (PI Helena Chui, MD), P30 AG066507 (PI Marilyn Albert, PhD), P30 AG066444 (PI David Holtzman, MD), P30 AG066518 (PI Lisa Silbert, MD, MCR), P30 AG066512 (PI Thomas Wisniewski, MD), P30 AG066462 (PI Scott Small, MD), P30 AG072979 (PI David Wolk, MD), P30 AG072972 (PI Charles DeCarli, MD), P30 AG072976 (PI Andrew Saykin, PsyD), P30 AG072975 (PI Julie A. Schneider, MD, MS), P30 AG072978 (PI Ann McKee, MD), P30 AG072977 (PI Robert Vassar, PhD), P30 AG066519 (PI Frank LaFerla, PhD), P30 AG062677 (PI Ronald Petersen, MD, PhD), P30 AG079280 (PI Jessica Langbaum, PhD), P30 AG062422 (PI Gil Rabinovici, MD), P30 AG066511 (PI Allan Levey, MD, PhD), P30 AG072946 (PI Linda Van Eldik, PhD), P30 AG062715 (PI Sanjay Asthana, MD, FRCP), P30 AG072973 (PI Russell Swerdlow, MD), P30 AG066506 (PI Glenn Smith, PhD, ABPP), P30 AG066508 (PI Stephen Strittmatter, MD, PhD), P30 AG066515 (PI Victor Henderson, MD, MS), P30 AG072947 (PI Suzanne Craft, PhD), P30 AG072931 (PI Henry Paulson, MD, PhD), P30 AG066546 (PI Sudha Seshadri, MD), P30 AG086401 (PI Erik Roberson, MD, PhD), P30 AG086404 (PI Gary Rosenberg, MD), P20 AG068082 (PI Angela Jefferson, PhD), P30 AG072958 (PI Heather Whitson, MD), P30 AG072959 (PI James Leverenz, MD).

Seonjoo Lee is partially supported by NIH R01AG062578.

