## Supplemental Table 1 to 11 for "Neuropathological Correlates of Apathy Progression in Alzheimer’s Disease and Related Dementias: A Longitudinal NACC Cohort Study"

### Supplementary eTable

eTable 1 Prevalence and Apathy Rates of the Neuropathological Combinations at Last Visit

| Neuropathology | cnt | Apathy cnt | Apathy Rate (%) |
| --- | --- | --- | --- |
| **AD+ CAA** | 238 | 113 | 47.5 |
| **AD+ LDB+ CAA** | 201 | 104 | 51.7 |
| **AD+ CAA+ CVD** | 120 | 47 | 39.2 |
| **NoNP** | 93 | 34 | 36.6 |
| **AD+ LDB+ CAA+ CVD** | 82 | 42 | 51.2 |
| **AD** | 73 | 33 | 45.2 |
| **FTLD** | 64 | 37 | 57.8 |
| **AD+ LDB** | 52 | 29 | 55.8 |
| **CVD** | 38 | 10 | 26.3 |
| **AD+ CAA+ HS** | 36 | 20 | 55.6 |
| **AD+ CVD** | 33 | 15 | 45.5 |
| **AD+ LDB+ CAA+ HS** | 30 | 18 | 60 |
| **CAA** | 27 | 7 | 25.9 |
| **LDB** | 26 | 13 | 50 |
| **AD+ LDB+ CAA+ CVD+ HS** | 22 | 11 | 50 |

**Notes**: Listed only the top 15 most common combinations. Abbreviations: AD, Alzheimer disease (defined as intermediate or high ABC criteria); CAA, cerebral amyloid angiopathy; CVD, cerebrovascular disease; FTLD, frontotemporal lobar degeneration; HS, hippocampal sclerosis; LBD, Lewy body disease; NoNP, no known pathology; cnt, count.

eTable 2 Association Between Apathy and Each Neuropathology Without Controlling for Other Pathologies Stratified by Sex

| ***Male*** | | | **OR (95% CI)** | | | | | | | | | | |
| --- | --- | --- | --- | --- | --- | --- | --- | --- | --- | --- | --- | --- | --- |
|  |  |  | **AD** | | **LBD** | | **CAA** | | **CVD** | | **FTLD** | | **HS** |
| **Model 1:** reference on absence of the target pathology | | **Pathology** | 1.03 | | **1.64*** | | 1 | | 1.12 | | **^+^2.58**** | | **^+^2.48**** |
|  |  |  | (0.62, 1.69) | | **(1.03, 2.61)** | | (0.62, 1.59) | | (0.68, 1.85) | | (1.40, 4.77) | | (1.29, 4.77) |
|  |  | **YTD** | **1.2***** | | **1.14***** | | **1.14***** | | **1.13***** | | **1.13***** | | **1.14***** |
|  |  |  | (1.10, 1.31) | | (1.08, 1.21) | | (1.06, 1.23) | | (1.07, 1.19) | | (1.06, 1.21) | | (1.08, 1.20) |
|  |  | **Pathology × YTD** | 0.96 | | 1.03 | | 1.02 | | 1.08 | | 1.01 | | 1.11 |
|  |  |  | (0.86, 1.06) | | (0.94, 1.13) | | (0.93, 1.12) | | (0.97, 1.19) | | (0.89, 1.15) | | (0.98, 1.27) |
| **Model 2:** reference on NoNP | | **Pathology** | 2.04 | | **2.67*** | | 2 | | 2.14 | | **3.63**** | | **4.31**** |
|  |  |  | (0.86, 4.84) | | (1.09, 6.52) | | (0.84, 4.76) | | (0.86, 5.37) | | (1.36, 9.72) | | (1.56, 11.94) |
|  |  | **YTD** | 1.07 | | 1.08 | | 1.07 | | 1.08 | | 1.09 | | 1.08 |
|  |  |  | (0.90, 1.27) | | (0.91, 1.28) | | (0.91, 1.28) | | (0.91, 1.28) | | (0.91, 1.30) | | (0.91, 1.28) |
|  |  | **Pathology × YTD** | 1.07 | | 1.09 | | 1.08 | | 1.13 | | 1.07 | | 1.18 |
|  |  |  | (0.89, 1.27) | | (0.91, 1.32) | | (0.91, 1.30) | | (0.93, 1.36) | | (0.87, 1.31) | | (0.96, 1.45) |
| ***Female*** | | | |  | |  | |  | |  |  |  | |
| **Model 1:** reference on absence of the target pathology | **Pathology** | | | 1.19 | | 1.16 | | 1.02 | | 0.94 | 1.59 | **2.18*** | |
|  |  |  |  | (0.63, 2.24) | | (0.65, 2.07) | | (0.58, 1.80) | | (0.52, 1.69) | (0.76, 3.35) | **(1.06, 4.47)** | |
|  | **YTD** | | | **1.14*** | | **1.13***** | | **1.16***** | | **1.14***** | **1.1*** | **1.14***** | |
|  |  |  |  | **(1.03, 1.27)** | | **(1.05, 1.20)** | | **(1.06, 1.26)** | | **(1.07, 1.21)** | **(1.02, 1.18)** | **(1.07, 1.22)** | |
|  | **Pathology × YTD** | | | 0.96 | | 0.99 | | 0.94 | | 0.93 | 1 | 0.94 | |
|  |  |  |  | (0.86, 1.08) | | (0.90, 1.09) | | (0.86, 1.04) | | (0.85, 1.03) | (0.87, 1.14) | (0.84, 1.06) | |
| **Model 2:** reference on NoNP | **Pathology** | | | 1.88 | | 2.03 | | 1.82 | | 1.78 | 1.89 | 3.48 | |
|  |  |  |  | (0.49, 7.27) | | (0.50, 8.24) | | (0.47, 7.10) | | (0.43, 7.34) | (0.42, 8.45) | (0.80, 15.20) | |
|  | **YTD** | | | 1.18 | | 1.18 | | 1.18 | | 1.17 | 1.2 | 1.19 | |
|  |  |  |  | (0.91, 1.52) | | (0.91, 1.53) | | (0.91, 1.52) | | (0.91, 1.52) | (0.92, 1.57) | (0.92, 1.54) | |
|  | **Pathology × YTD** | | | 0.94 | | 0.94 | | 0.93 | | 0.91 | 0.9 | 0.91 | |
|  |  |  |  | (0.73, 1.21) | | (0.72, 1.22) | | (0.72, 1.20) | | (0.70, 1.18) | (0.68, 1.21) | (0.69, 1.19) | |

**Notes:** Both models adjusted for sex, age at death, and NPI-Q total score excluding the apathy item. Significance level, P ≤ 0.05*, P ≤ 0.01**, P ≤ 0.001***, **^+^** P < .05 after multiple comparisons correction to control for false discovery rate. Abbreviations: AD, Alzheimer disease (defined as intermediate or high ABC criteria); CAA, cerebral amyloid angiopathy; CVD, cerebrovascular disease; FTLD, frontotemporal lobar degeneration; HS, hippocampal sclerosis; LBD, Lewy body disease; NoNP, no known pathology; OR, odds ratio; CI, confidence interval; YTD, year to death.

eTable 3 Association Between Apathy Severity and Each Neuropathology Without Controlling for Other Pathologies

|  | | **OR (95% CI)** | | | | | |
| --- | --- | --- | --- | --- | --- | --- | --- |
|  |  | **AD** | **LBD** | **CAA** | **CVD** | **FTLD** | **HS** |
| **Model 1:** reference on absence of the target pathology | **Pathology** | 1.1 | **1.44*** | 1.14 | 0.94 | **^+^2.32***** | **^+^2.29***** |
|  |  | (0.77, 1.58) | (1.04, 2.00) | (0.82, 1.59) | (0.66, 1.34) | (1.55, 3.46) | (1.47, 3.56) |
|  | **YTD** | **1.19***** | **1.15***** | **1.15***** | **1.16***** | **1.12***** | **1.16***** |
|  |  | (1.12, 1.26) | (1.11, 1.20) | (1.12, 1.21) | (1.12, 1.21) | (1.08, 1.17) | (1.12, 1.20) |
|  | **Pathology × YTD** | 0.97 | 1.02 | 1 | 0.98 | 1.05 | 1.04 |
|  |  | (0.90, 1.03) | (0.96, 1.08) | (0.94, 1.07) | (0.92, 1.05) | (0.98, 1.14) | (0.96, 1.12) |
| **Model 2:** reference on NoNP | **Pathology** | 1.61 | 1.93 | 1.63 | 1.49 | **2.32*** | **^+^3.17**** |
|  |  | (0.82, 3.15) | (0.97, 3.86) | (0.83, 3.21) | (0.73, 3.04) | (1.13, 4.75) | (1.47, 6.81) |
|  | **YTD** | 1.16 | 1.16 | 1.16 | 1.16 | 1.17 | 1.16 |
|  |  | (1.02, 1.31) | (1.02, 1.32) | (1.02, 1.32) | (1.02, 1.32) | (1.03, 1.33) | (1.02, 1.32) |
|  | **Pathology × YTD** | 0.99 | 1.01 | 1 | 0.99 | 1 | 1.03 |
|  |  | (0.87, 1.13) | (0.88, 1.15) | (0.88, 1.14) | (0.86, 1.13) | (0.87, 1.16) | (0.89, 1.19) |

**Notes:** Both models adjusted for sex, age at death, and NPI-Q total score excluding the apathy item. Significance level, P ≤ 0.05*, P ≤ 0.01**, P ≤ 0.001***, **^+^** P < .05 after multiple comparisons correction to control for false discovery rate. Abbreviations: AD, Alzheimer disease (defined as intermediate or high ABC criteria); CAA, cerebral amyloid angiopathy; CVD, cerebrovascular disease; FTLD, frontotemporal lobar degeneration; HS, hippocampal sclerosis; LBD, Lewy body disease; NoNP, no known pathology; OR, odds ratio; CI, confidence interval; YTD, year to death.

eTable 4 Association Between Apathy Severity and Each Neuropathology Without Controlling for Other Pathologies, Stratified by Sex

| ***Male*** | | | **OR (95% CI)** | | | | | | | | | | |
| --- | --- | --- | --- | --- | --- | --- | --- | --- | --- | --- | --- | --- | --- |
|  |  |  | **AD** | | **LBD** | | **CAA** | | **CVD** | | **FTLD** | | **HS** |
| **Model 1:** reference on absence of the target pathology | | **Pathology** | 1.11 | | 1.46 | | 1.18 | | 0.98 | | **^+^2.39***** | | **^+^2.26**** |
|  |  |  | (0.71, 1.74) | | (0.96, 2.21) | | (0.77, 1.79) | | (0.62, 1.54) | | (1.43, 4.01) | | (1.25, 4.06) |
|  |  | **YTD** | **1.19***** | | **1.16***** | | **1.15***** | | **1.16***** | | **1.14***** | | **1.16***** |
|  |  |  | (1.11, 1.29) | | (1.10, 1.23) | | (1.11, 1.23) | | (1.11, 1.23) | | (1.08, 1.21) | | (1.11, 1.22) |
|  |  | **Pathology × YTD** | 0.98 | | 1.03 | | 1.04 | | 1.02 | | 1.03 | | 1.11 |
|  |  |  | (0.89, 1.07) | | (0.95, 1.11) | | (0.96, 1.13) | | (0.94, 1.12) | | (0.93, 1.15) | | (0.99, 1.25) |
| **Model 2:** reference on NoNP | | **Pathology** | 2.04 | | **2.67*** | | 2 | | 2.14 | | **3.63**** | | **4.31**** |
|  |  |  | (0.86, 4.84) | | (1.09, 6.52) | | (0.84, 4.76) | | (0.86, 5.37) | | (1.36, 9.72) | | (1.56, 11.94) |
|  |  | **YTD** | 1.07 | | 1.08 | | 1.07 | | 1.08 | | 1.09 | | 1.08 |
|  |  |  | (0.90, 1.27) | | (0.91, 1.28) | | (0.91, 1.28) | | (0.91, 1.28) | | (0.91, 1.30) | | (0.91, 1.28) |
|  |  | **Pathology × YTD** | 1.07 | | 1.09 | | 1.08 | | 1.13 | | 1.07 | | 1.18 |
|  |  |  | (0.89, 1.27) | | (0.91, 1.32) | | (0.91, 1.30) | | (0.93, 1.36) | | (0.87, 1.31) | | (0.96, 1.45) |
| ***Female*** | | | |  | |  | |  | |  |  |  | |
| **Model 1:** reference on absence of the target pathology | **Pathology** | | | 1.09 | | 1.42 | | 1.1 | | 0.91 | **2.17*** | **2.45**** | |
|  |  |  |  | (0.60, 2.01) | | (0.83, 2.43) | | (0.64, 1.88) | | (0.52, 1.59) | (1.13, 4.14) | (1.25, 4.83) | |
|  | **YTD** | | | **1.17*** | | **1.13***** | | **1.16***** | | **1.16***** | **1.1*** | **1.15***** | |
|  |  |  |  | (1.05, 1.29) | | (1.06, 1.21) | | (1.09, 1.23) | | (1.09, 1.23) | (1.03, 1.18) | (1.08, 1.22) | |
|  | **Pathology × YTD** | | | 0.96 | | 1.01 | | 0.96 | | 0.94 | 1.08 | 0.98 | |
|  |  |  |  | (0.86, 1.06) | | (0.93, 1.11) | | (0.87, 1.05) | | (0.85, 1.02) | (0.96, 1.22) | (0.88, 1.09) | |
| **Model 2:** reference on NoNP | **Pathology** | | | 1.88 | | 2.03 | | 1.82 | | 1.78 | 1.89 | 3.48 | |
|  |  |  |  | (0.49, 7.27) | | (0.50, 8.24) | | (0.47, 7.10) | | (0.43, 7.34) | (0.42, 8.45) | (0.80, 15.20) | |
|  | **YTD** | | | 1.18 | | 1.18 | | 1.18 | | 1.17 | 1.2 | 1.19 | |
|  |  |  |  | (0.91, 1.52) | | (0.91, 1.53) | | (0.91, 1.52) | | (0.91, 1.52) | (0.92, 1.57) | (0.92, 1.54) | |
|  | **Pathology × YTD** | | | 0.94 | | 0.94 | | 0.93 | | 0.91 | 0.9 | 0.91 | |
|  |  |  |  | (0.73, 1.21) | | (0.72, 1.22) | | (0.72, 1.20) | | (0.70, 1.18) | (0.68, 1.21) | (0.69, 1.19) | |

**Notes:** Both models adjusted for sex, age at death, and NPI-Q total score excluding the apathy item. Significance level, P ≤ 0.05*, P ≤ 0.01**, P ≤ 0.001***, **^+^** P < .05 after multiple comparisons correction to control for false discovery rate. Abbreviations: AD, Alzheimer disease (defined as intermediate or high ABC criteria); CAA, cerebral amyloid angiopathy; CVD, cerebrovascular disease; FTLD, frontotemporal lobar degeneration; HS, hippocampal sclerosis; LBD, Lewy body disease; NoNP, no known pathology; OR, odds ratio; CI, confidence interval; YTD, year to death.

eTable 5 Independent Effect of Neuropathologies on Apathy Progression Stratified by Sex

| Terms | Male | | | | Female | | | |
| --- | --- | --- | --- | --- | --- | --- | --- | --- |
|  | **Model 3** | | **Model 4** | | **Model 3** | | **Model 4** | |
|  | **OR** | **95% CI** | **OR** | **95% CI** | **OR** | **95% CI** | **OR** | **95% CI** |
| **YTD** | 1.12 | (0.98, 1.28) | 1.16 | (0.99, 1.36) | **1.24**** | (1.06, 1.45) | **1.24*** | (1.04, 1.47) |
| **AD** | 1.18 | (0.57, 2.42) | 1.09 | (0.51, 2.34) | 1.04 | (0.43, 2.52) | 1.04 | (0.41, 2.64) |
| **LBD** | 1.77 | (0.99, 3.16) | 1.73 | (0.96, 3.10) | 1.1 | (0.55, 2.21) | 1.1 | (0.55, 2.22) |
| **CAA** | 0.9 | (0.46, 1.73) | 0.87 | (0.45, 1.69) | 1.13 | (0.54, 2.38) | 1.13 | (0.53, 2.40) |
| **CVD** | 1.29 | (0.71, 2.34) | 1.25 | (0.68, 2.29) | 0.75 | (0.38, 1.50) | 0.75 | (0.37, 1.52) |
| **FTLD** | **3.28***** | (1.63, 6.60) | **3.16***** | (1.56, 6.40) | 1.42 | (0.60, 3.34) | 1.42 | (0.60, 3.36) |
| **HS** | **2.8**** | (1.31, 5.97) | **2.72**** | (1.27, 5.84) | 2.05 | (0.86, 4.86) | 2.05 | (0.86, 4.88) |
| **NoNP** |  |  | 0.71 | (0.23, 2.21) |  |  | 1.04 | (0.21, 5.23) |
| **AD × YTD** | 0.91 | (0.79, 1.06) | 0.9 | (0.77, 1.05) | 0.92 | (0.78, 1.08) | 0.92 | (0.78, 1.09) |
| **LBD × YTD** | 1.07 | (0.95, 1.20) | 1.06 | (0.94, 1.19) | 1.02 | (0.90, 1.14) | 1.02 | (0.90, 1.15) |
| **CAA × YTD** | 1 | (0.87, 1.14) | 0.99 | (0.87, 1.13) | 1.03 | (0.90, 1.17) | 1.03 | (0.90, 1.17) |
| **CVD × YTD** | 1.1 | (0.98, 1.24) | 1.09 | (0.97, 1.23) | 0.89 | (0.79, 1.00) | 0.89 | (0.79, 1.00) |
| **FTLD × YTD** | 1.01 | (0.88, 1.17) | 1 | (0.87, 1.16) | 0.94 | (0.81, 1.10) | 0.94 | (0.81, 1.10) |
| **HS × YTD** | 1.16 | (1.00, 1.35) | 1.16 | (0.99, 1.34) | 0.94 | (0.82, 1.08) | 0.94 | (0.82, 1.09) |
| **NoNP × YTD** |  |  | 0.92 | (0.72, 1.17) |  |  | 1.02 | (0.75, 1.39) |

**Notes:** Both models assessed the independent effect of all neuropathologies simultaneously on apathy progression, while controlling for sex, age at death, and NPI-Q total score excluding the apathy item. Model 4 additionally adjusted for NP. Significance level, P ≤ 0.05*, P ≤ 0.01**, P ≤ 0.001***. Abbreviations: AD, Alzheimer disease (defined as intermediate or high ABC criteria); CAA, cerebral amyloid angiopathy; CVD, cerebrovascular disease; FTLD, frontotemporal lobar degeneration; HS, hippocampal sclerosis; LBD, Lewy body disease; NoNP, no known pathology; OR, odds ratio; CI, confidence interval; YTD, year to death.

eTable 6 Association Between Apathy and Comorbidity Without Controlling for Other Pathologies

|  |  | OR (95% CI) | | | |
| --- | --- | --- | --- | --- | --- |
|  |  | **AD+CAA** | **AD+LBD+CAA** | **AD+CAA+CVD** | |
| **Model 5:** reference on absence of the target pathology | **Pathology** | 0.84 | 0.93 | | 0.79 |
|  |  | (0.52, 1.34) | (0.55, 1.55) | | (0.41, 1.50) |
|  | **YTD** | 1.13*** | 1.14*** | | 1.15*** |
|  |  | (1.09, 1.17) | (1.10, 1.19) | | (1.11, 1.19) |
|  | **Pathology:YTD** | 1.06 | 0.98 | | 0.91 |
|  |  | (0.97, 1.15) | (0.89, 1.07) | | (0.82, 1.02) |
| **Model 6:** reference on NoNP | **Pathology** | 1.7 | 1.83 | | 1.58 |
|  |  | (0.75, 3.86) | (0.79, 4.26) | | (0.62, 4.03) |
|  | **YTD** | 1.1 | 1.1 | | 1.1 |
|  |  | (0.96, 1.26) | (0.96, 1.26) | | (0.96, 1.26) |
|  | **Pathology:YTD** | 1.09 | 1.02 | | 0.95 |
|  |  | (0.93, 1.28) | (0.86, 1.20) | | (0.80, 1.14) |

**Notes**: Listed only the top 3 most common combinations. Abbreviations: AD, Alzheimer disease (defined as intermediate or high ABC criteria); CAA, cerebral amyloid angiopathy; CVD, cerebrovascular disease; LBD, Lewy body disease; NoNP, no known pathology.

eTable 7 Association Between Apathy and Each Neuropathology Without Controlling for Other Pathologies, Additionally Adjusted for Depression

|  |  | **OR (95% CI)** | | | | | |
| --- | --- | --- | --- | --- | --- | --- | --- |
|  |  | **AD** | **LBD** | **CAA** | **CVD** | **FTLD** | **HS** |
| **Model 1:** reference on absence of the target pathology | **Pathology** | 1.09 | 1.41 | 1.01 | 0.99 | **^+^2.23***** | **^+^2.35***** |
|  |  | (0.74, 1.61) | (0.98, 2.02) | (0.71, 1.45) | (0.68, 1.45) | (1.39, 3.58) | (1.46, 3.80) |
|  | **YTD** | **1.19***** | **1.15***** | **1.16***** | **1.15***** | **1.13***** | **1.16***** |
|  |  | (1.11, 1.27) | (1.10, 1.20) | (1.10, 1.20) | (1.10, 1.20) | (1.07, 1.18) | (1.11, 1.20) |
|  | **Pathology × YTD** | 0.96 | 1.01 | 0.98 | 1 | 1.01 | 1.01 |
|  |  | (0.89, 1.03) | (0.94, 1.08) | (0.92, 1.05) | (0.94, 1.07) | (0.93, 1.11) | (0.93, 1.10) |
| **Model 2:** reference on NoNP | **Pathology** | 1.93 | **2.33*** | 1.88 | 1.87 | **2.8*** | **^+^3.88***** |
|  |  | (0.93, 4.01) | (1.09, 4.96) | (0.90, 3.92) | (0.87, 4.02) | (1.23, 6.37) | (1.69, 8.88) |
|  | **YTD** | 1.12 | 1.12 | 1.12 | 1.12 | 1.13 | 1.13 |
|  |  | (0.97, 1.28) | (0.98, 1.29) | (0.98, 1.29) | (0.98, 1.29) | (0.98, 1.31) | (0.98, 1.30) |
|  | **Pathology × YTD** | 1.02 | 1.03 | 1.02 | 1.03 | 1.01 | 1.04 |
|  |  | (0.88, 1.17) | (0.89, 1.20) | (0.88, 1.18) | (0.88, 1.19) | (0.85, 1.19) | (0.89, 1.22) |

**Notes:** Both models adjusted for sex, age at death, depression, and NPI-Q total score excluding the apathy item. Significance level, P ≤ 0.05*, P ≤ 0.01**, P ≤ 0.001***, **^+^** P < .05 after multiple comparisons correction to control for false discovery rate. Abbreviations: AD, Alzheimer disease (defined as intermediate or high ABC criteria); CAA, cerebral amyloid angiopathy; CVD, cerebrovascular disease; FTLD, frontotemporal lobar degeneration; HS, hippocampal sclerosis; LBD, Lewy body disease; NoNP, no known pathology; OR, odds ratio; CI, confidence interval; YTD, year to death.

eTable 8 Association Between Apathy and Each Neuropathology Without Controlling for Other Pathologies, Stratified by Sex, Additionally Adjusted for Depression

| ***Male*** | | **OR (95% CI)** | | | | | | | | | |
| --- | --- | --- | --- | --- | --- | --- | --- | --- | --- | --- | --- |
|  |  | **AD** | | **LBD** | | **CAA** | | **CVD** | | **FTLD** | **HS** |
| **Model 1:** reference on absence of the target pathology | **Pathology** | 1.05 | | **1.6*** | | 1.01 | | 1.09 | | **^+^2.74**** | **^+^2.66***** |
|  |  | (0.64, 1.72) | | (1.01, 2.56) | | (0.63, 1.62) | | (0.66, 1.79) | | (1.49, 5.06) | (1.39, 5.10) |
|  | **YTD** | **1.2***** | | **1.15***** | | **1.15***** | | **1.14***** | | **1.14***** | **1.15***** |
|  |  | (1.10, 1.31) | | (1.08, 1.22) | | (1.08, 1.20) | | (1.08, 1.20) | | (1.06, 1.21) | (1.09, 1.21) |
|  | **Pathology × YTD** | 0.96 | | 1.03 | | 1.02 | | 1.07 | | 1.02 | 1.12 |
|  |  | (0.86, 1.06) | | (0.94, 1.13) | | (0.93, 1.12) | | (0.97, 1.18) | | (0.90, 1.16) | (0.99, 1.28) |
| **Model 2:** reference on NoNP | **Pathology** | 1.95 | | **2.51*** | | 1.92 | | 2 | | **3.51*** | **4.36**** |
|  |  | (0.82, 4.64) | | (1.03, 6.15) | | (0.80, 4.57) | | (0.80, 5.01) | | (1.31, 9.39) | (1.57, 12.07) |
|  | **YTD** | 1.09 | | 1.1 | | 1.09 | | 1.1 | | 1.11 | 1.1 |
|  |  | (0.92, 1.29) | | (0.92, 1.30) | | (0.93, 1.30) | | (0.93, 1.30) | | (0.93, 1.32) | (0.92, 1.30) |
|  | **Pathology × YTD** | 1.05 | | 1.08 | | 1.07 | | 1.11 | | 1.06 | 1.17 |
|  |  | (0.88, 1.26) | | (0.90, 1.30) | | (0.90, 1.28) | | (0.92, 1.34) | | (0.86, 1.30) | (0.96, 1.44) |
| ***Female*** | | |  | |  | |  | |  |  |  |
| **Model 1:** reference on absence of the target pathology | **Pathology** | | 1.2 | | 1.16 | | 1.04 | | 0.93 | 1.67 | **2.28*** |
|  |  |  | (0.63, 2.26) | | (0.65, 2.07) | | (0.59, 1.82) | | (0.52, 1.68) | (0.80, 3.51) | (1.11, 4.68) |
|  | **YTD** | | **1.15**** | | **1.13***** | | **1.17***** | | **1.15***** | **1.1*** | **1.15***** |
|  |  |  | (1.04, 1.28) | | (1.06, 1.21) | | (1.07, 1.22) | | (1.07, 1.22) | (1.02, 1.19) | (1.08, 1.23) |
|  | **Pathology × YTD** | | 0.96 | | 0.99 | | 0.94 | | 0.93 | 1 | 0.94 |
|  |  |  | (0.86, 1.08) | | (0.90, 1.09) | | (0.85, 1.04) | | (0.85, 1.03) | (0.87, 1.15) | (0.84, 1.06) |
| **Model 2:** reference on NoNP | **Pathology** | | 1.85 | | 1.98 | | 1.79 | | 1.73 | 1.86 | 3.53 |
|  |  |  | (0.48, 7.16) | | (0.49, 8.10) | | (0.46, 7.00) | | (0.42, 7.16) | (0.42, 8.31) | (0.81, 15.43) |
|  | **YTD** | | 1.2 | | 1.2 | | 1.19 | | 1.19 | 1.22 | 1.21 |
|  |  |  | (0.93, 1.55) | | (0.93, 1.56) | | (0.92, 1.54) | | (0.92, 1.54) | (0.93, 1.60) | (0.93, 1.57) |
|  | **Pathology × YTD** | | 0.93 | | 0.93 | | 0.92 | | 0.9 | 0.89 | 0.9 |
|  |  |  | (0.72, 1.20) | | (0.71, 1.21) | | (0.71, 1.19) | | (0.69, 1.17) | (0.67, 1.20) | (0.68, 1.18) |

**Notes:** Both models adjusted for sex, age at death, depression, and NPI-Q total score excluding the apathy item. Significance level, P ≤ 0.05*, P ≤ 0.01**, P ≤ 0.001***, **^+^** P < .05 after multiple comparisons correction to control for false discovery rate. Abbreviations: AD, Alzheimer disease (defined as intermediate or high ABC criteria); CAA, cerebral amyloid angiopathy; CVD, cerebrovascular disease; FTLD, frontotemporal lobar degeneration; HS, hippocampal sclerosis; LBD, Lewy body disease; NoNP, no known pathology; OR, odds ratio; CI, confidence interval; YTD, year to death.

eTable 9 Independent Effect of Neuropathologies on Apathy Progression, Additionally Adjust for Depression

| **Terms** | **Model 3** | | **Model 4** | |
| --- | --- | --- | --- | --- |
|  | **OR** | **95% CI** | **OR** | **95% CI** |
| **YTD** | **1.2***** | (1.08, 1.33) | **1.22***** | (1.08, 1.37) |
| **AD** | 1.12 | (0.64, 1.96) | 1.07 | (0.59, 1.94) |
| **LBD** | 1.4 | (0.90, 2.19) | 1.39 | (0.88, 2.17) |
| **CAA** | 0.99 | (0.60, 1.62) | 0.98 | (0.59, 1.61) |
| **CVD** | 0.94 | (0.60, 1.48) | 0.93 | (0.59, 1.47) |
| **FTLD** | **2.36**** | (1.38, 4.06) | **2.32**** | (1.35, 4.01) |
| **HS** | **2.36**** | (1.34, 4.17) | **2.34**** | (1.32, 4.13) |
| **NoNP** |  |  | 0.82 | (0.32, 2.09) |
| **AD × YTD** | 0.92 | (0.82, 1.02) | 0.91 | (0.81, 1.02) |
| **LBD × YTD** | 1.03 | (0.95, 1.12) | 1.03 | (0.95, 1.12) |
| **CAA × YTD** | 1.01 | (0.92, 1.11) | 1 | (0.91, 1.10) |
| **CVD × YTD** | 0.99 | (0.91, 1.08) | 0.99 | (0.91, 1.08) |
| **FTLD × YTD** | 0.98 | (0.88, 1.09) | 0.97 | (0.88, 1.08) |
| **HS × YTD** | 1.03 | (0.93, 1.14) | 1.03 | (0.93, 1.14) |
| **NoNP × YTD** |  |  | 0.95 | (0.78, 1.15) |

**Notes:** Both models assessed the independent effect of all neuropathologies simultaneously on apathy progression, while controlling for sex, age at death, depression and NPI-Q total score excluding the apathy item. Model 4 additionally adjusted for NP. Significance level, P ≤ 0.05*, P ≤ 0.01**, P ≤ 0.001***. Abbreviations: AD, Alzheimer disease (defined as intermediate or high ABC criteria); CAA, cerebral amyloid angiopathy; CVD, cerebrovascular disease; FTLD, frontotemporal lobar degeneration; HS, hippocampal sclerosis; LBD, Lewy body disease; NoNP, no known pathology; OR, odds ratio; CI, confidence interval; YTD, year to death.

eTable 10 Independent Effect of Neuropathologies on Apathy Progression Stratified by Sex, Additionally Adjust for Depression

| Terms | Male | | | | Female | | | |
| --- | --- | --- | --- | --- | --- | --- | --- | --- |
|  | **Model 3** | | **Model 4** | | **Model 3** | | **Model 4** | |
|  | **OR** | **95% CI** | **OR** | **95% CI** | **OR** | **95% CI** | **OR** | **95% CI** |
| **YTD** | 1.13 | (0.99, 1.29) | 1.15 | (0.98, 1.35) | **1.25**** | (1.07, 1.47) | **1.25*** | (1.05, 1.48) |
| **AD** | 1.18 | (0.57, 2.44) | 1.14 | (0.53, 2.45) | 1.04 | (0.43, 2.52) | 1.05 | (0.42, 2.66) |
| **LBD** | 1.74 | (0.98, 3.11) | 1.72 | (0.96, 3.09) | 1.09 | (0.54, 2.20) | 1.1 | (0.54, 2.21) |
| **CAA** | 0.91 | (0.47, 1.75) | 0.9 | (0.46, 1.74) | 1.15 | (0.54, 2.42) | 1.15 | (0.54, 2.44) |
| **CVD** | 1.26 | (0.70, 2.28) | 1.25 | (0.68, 2.28) | 0.74 | (0.37, 1.48) | 0.75 | (0.37, 1.50) |
| **FTLD** | **3.49***** | (1.73, 7.01) | **3.42***** | (1.68, 6.94) | 1.49 | (0.63, 3.51) | 1.5 | (0.63, 3.54) |
| **HS** | **3.07**** | (1.44, 6.54) | **3.04**** | (1.42, 6.51) | 2.13 | (0.90, 5.06) | 2.14 | (0.90, 5.09) |
| **NoNP** |  |  | 0.86 | (0.27, 2.70) |  |  | 1.1 | (0.22, 5.48) |
| **AD × YTD** | 0.91 | (0.79, 1.06) | 0.9 | (0.77, 1.06) | 0.92 | (0.78, 1.08) | 0.92 | (0.78, 1.09) |
| **LBD × YTD** | 1.07 | (0.95, 1.20) | 1.06 | (0.95, 1.20) | 1.01 | (0.90, 1.14) | 1.01 | (0.90, 1.14) |
| **CAA × YTD** | 1 | (0.88, 1.14) | 1 | (0.87, 1.14) | 1.03 | (0.90, 1.17) | 1.03 | (0.90, 1.17) |
| **CVD × YTD** | 1.1 | (0.98, 1.24) | 1.09 | (0.97, 1.23) | 0.89 | (0.79, 1.00) | 0.89 | (0.79, 1.00) |
| **FTLD × YTD** | 1.02 | (0.89, 1.18) | 1.01 | (0.88, 1.17) | 0.95 | (0.81, 1.11) | 0.95 | (0.81, 1.11) |
| **HS × YTD** | 1.17 | (1.01, 1.36) | 1.17 | (1.01, 1.36) | 0.94 | (0.82, 1.08) | 0.94 | (0.82, 1.09) |
| **NoNP × YTD** |  |  | 0.95 | (0.74, 1.21) |  |  | 1.03 | (0.75, 1.41) |

**Notes:** Both models assessed the independent effect of all neuropathologies simultaneously on apathy progression, while controlling for sex, age at death, depression and NPI-Q total score excluding the apathy item. Model 4 additionally adjusted for NP. Significance level, P ≤ 0.05*, P ≤ 0.01**, P ≤ 0.001***. Abbreviations: AD, Alzheimer disease (defined as intermediate or high ABC criteria); CAA, cerebral amyloid angiopathy; CVD, cerebrovascular disease; FTLD, frontotemporal lobar degeneration; HS, hippocampal sclerosis; LBD, Lewy body disease; NoNP, no known pathology; OR, odds ratio; CI, confidence interval; YTD, year to death.

eTable 11 Distribution of FTLD Subtypes

| **FTLD subtypes** | **cnt** | **Apathy cnt** | **Apathy Rate(%)** |
| --- | --- | --- | --- |
| **Other Pathologies** | 858 | 370 | 43.1 |
| **Missing (in FTLD)** | 334 | 171 | 51.2 |
| **TAU** | 205 | 105 | 51.2 |
| **TDP-43** | 68 | 48 | 70.6 |
| **TDP-43 + TAU** | 20 | 15 | 75 |
| **Other** | 1 | 0 | 0 |
| **Other + TDP-43** | 1 | 1 | 100 |
| **Other + TDP-43 + TAU** | 1 | 1 | 100 |

**Notes**: ‘Other’ includes FTLD-FUS and others. Abbreviations: FTLD, frontotemporal lobar degeneration.
