## Supplemental Figure 1 for "Neuropathological Correlates of Apathy Progression in Alzheimer’s Disease and Related Dementias: A Longitudinal NACC Cohort Study"

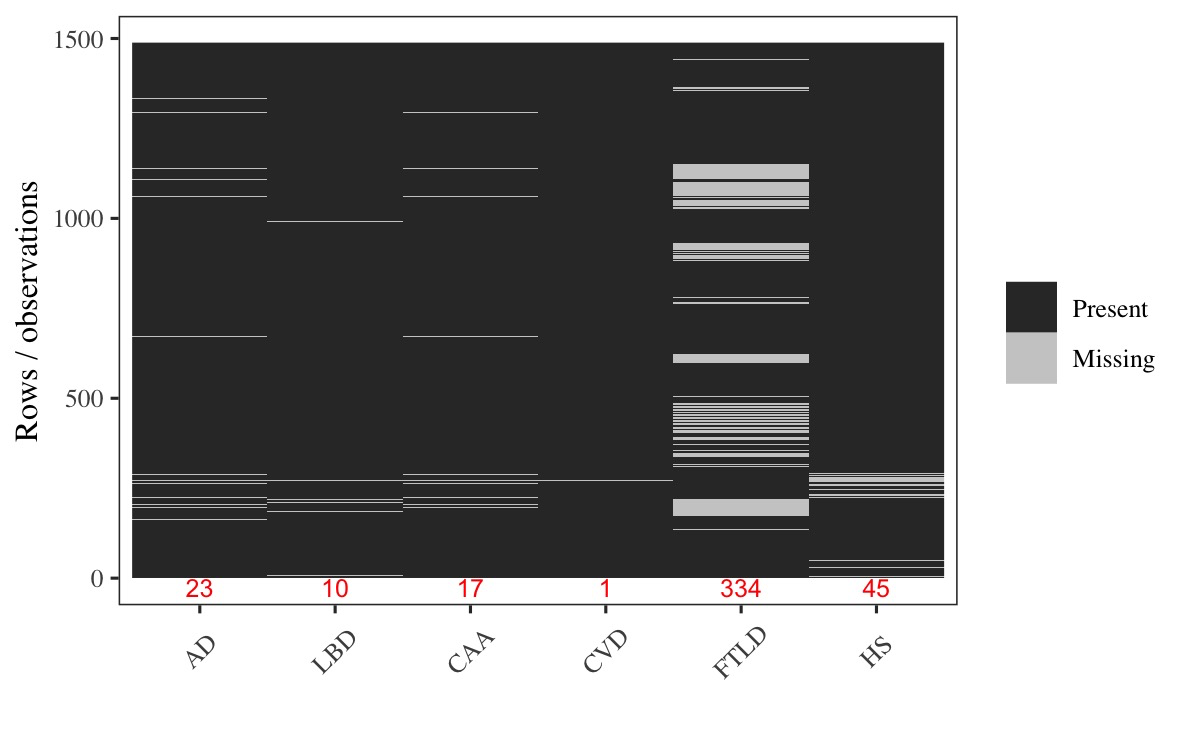

### Supplementary eFigure

eFigure 1 Missing data pattern for neuropathological variables. Each row represents an individual observation; each column represents one pathology: Alzheimer’s disease (AD), Lewy body disease (LBD), cerebral amyloid angiopathy (CAA), cerebrovascular disease (CVD), frontotemporal lobar degeneration (FTLD), and hippocampal sclerosis (HS). Black indicates available data, and gray indicates missing data. Red numbers at the bottom denote the count of missing observations for each variable.
